# Peptidomimetic inhibitors targeting TrkB/PSD-95 signaling improves cognition and seizure outcomes in an Angelman Syndrome mouse model

**DOI:** 10.1101/2024.06.07.597833

**Authors:** Emily Z. Huie, Xin Yang, Mengia S. Rioult-Pedotti, Mandar Naik, Yu-Wen Alvin Huang, Jill L Silverman, John Marshall

## Abstract

Angelman syndrome (AS) is a rare genetic neurodevelopmental disorder with profoundly debilitating symptoms with no FDA-approved cure or therapeutic. Brain-derived neurotrophic factor (BDNF), and its receptor TrkB, have a well-established role as regulators of synaptic plasticity, dendritic outgrowth, dendritic spine formation and maintenance. Previously, we reported that the association of PSD-95 with TrkB is critical for intact BDNF signaling in the AS mouse model, as illustrated by attenuated PLCγ and PI3K signaling and intact MAPK pathway signaling. These data suggest that drugs tailored to enhance the TrkB-PSD-95 interaction may provide a novel approach for the treatment of AS and a variety of NDDs. To evaluate this critical interaction, we synthesized a class of high-affinity PSD-95 ligands that bind specifically to the PDZ3 domain of PSD-95, denoted as Syn3 peptidomimetic ligands. We evaluated Syn3 and its analog D-Syn3 (engineered using dextrorotary (D)-amino acids) *in vivo* using the *Ube3a* exon 2 deletion mouse model of AS. Following systemic administration of Syn3 and D-Syn3, we demonstrated improvement in the seizure domain of AS. Learning and memory using the novel object recognition assay also illustrated improved cognition following Syn3 and D-Syn3, along with restored long-term potentiation. Finally, D-Syn3 treated mice showed a partial rescue in motor learning. Neither Syn3 nor D-Syn3 improved gross exploratory locomotion deficits, nor gait impairments that have been well documented in the AS rodent models. These findings highlight the need for further investigation of this compound class as a potential therapeutic for AS and other genetic NDDs.

## Introduction

Angelman syndrome (AS) is a rare genetic neurodevelopmental disorder (NDD) with debilitating symptoms, including gross and fine motor impairments, poor motor coordination, gait deficits, intellectual disability, and recurrent, uncontrollable seizures [1–4]. Symptoms of AS are severe and lifelong, yet there are no FDA-approved therapeutics that address the major symptom domains of this disorder . The loss of expression in the maternal allele of ubiquitin ligase E3A (*UBE3A)* gene results in a myriad of downstream effects, including affecting synaptic proteins that are theorized to be contributors to many of the phenotypes observed in AS. Animal models are essential for evaluating the efficacy and safety of potential therapeutics *in vivo.* The *Ube3a* exon 2 mouse model successfully recapitulates translationally relevant behaviors including, poor motor abilities and coordination, abnormal gait, cognitive deficits, and heightened seizure susceptibility, thus serves as a reliable *in vivo* model to test potential AS therapeutics [5–11].

Current AS therapeutic endeavors range from genetic precision medicine to traditional drug therapies, yet delivery to the brain remains challenging for both routes. The consistent finding of abnormal synaptic dendritic spine morphology, decreased dendritic spine density, and reduced long-term potentiation (LTP) in AS mice [12–16] led us to explore whether deficits in brain-derived neurotrophic factor (BDNF) signaling, which has well-established roles in the regulation of synaptic plasticity as well dendritic spine formation, growth, and maintenance [17–29], could underlie AS pathophysiology. As reported, the association between TrkB and PSD-95 was found to be critical for intact BDNF signaling [30], and this association was disrupted in AS mice resulting in attenuated PLCγ-CaMKII and PI3K signaling [30,31]. These data suggest that drugs tailored to enhance TrkB-PSD-95 interaction may be a novel approach for the treatment of AS and a variety of NDDs.

To evaluate this interaction, generate, and optimize a lead candidate, we synthesized a class of high-affinity PSD-95 peptidomimetic ligands that bind to the PDZ3-domain of PSD-95. Using structure-based knowledge, we developed new peptidomimetic compounds, Syn3 and D-Syn3 (an analog engineered using dextrorotary (D)-amino acids), referred to as Syn3 compounds, that fuse peptides derived from the PSD-95-binding protein SynGAP1 to our prototype compound CN2097 [30,32]. The new compounds target both the 3^rd^ PDZ subunit of PSD-95 and adjoining αC helix to achieve bivalent binding that results in up to 7-fold stronger affinity compared to CN2097 [32].

Given the novelty of the Syn3 compounds and their promotion of BDNF-TrkB signaling at synapses, we performed behavioral and seizure assays, tailored to AS, to examine the efficacy of two Syn compounds *in vivo* in AS and WT littermate control mice [8]. Our objectives in this report were 1) to assess the preliminary active dose of peptidomimetics, Syn3 and D-Syn3, and their postulated mechanism of action, as novel therapeutics for AS; 2) identify improved symptom domain(s) in the AS model, providing a path forward for discovery and assessment in other genetic NDDs, which share phenotypic features, and may also benefit from these compounds. Our data indicate this compound class is efficacious for the symptom domains of learning and memory, seizures, underlying LTP, and thus E/I imbalance overall.

## Materials and Methods

### Peptide synthesis of Syn3 and D-Syn3

Syn3 and D-Syn3, bridged cyclic peptidomimetic compounds were synthesized by WuXi AppTec using 9-fluorenylmethoxycarbonyl (Fmoc) solid phase peptide synthesis protocols, as previously published [32]. Compounds were purified by RP-HPLC, lyophilized, and exchanged with HCl. Peptide purity was greater than 95% as determined using high-resolution time of flight on an AXIMA Performance MALDI TOF/TOF mass spectrometer (Shimadzu; Kyoto, Japan).

### Electrophysiology in hippocampal slices to assess deficits in long term potentiation in AS mice and the effects of Syn3 and D-Syn3

Slice preparation, electrophysiological recordings, and data analysis were performed as described and are further detailed in the Supplementary Materials [31].

### Pluripotent stem cell induced neurons (iNs)

iNs were generated following previous protocol by forced expression of Neurogenin-2 (NGN2) with kolf2.J1 stem cell lines [33,34]. Details of the model system and our assay work are further described in the Supplementary Materials.

### Western blot

All samples were harvested in cell lysis buffer (1% SDS, 0.5% Deoxcholate, 50 mM Sodium phosphate diabasic heptahydrate, 150 mM Sodium chloride, 2mM EDTA, 50 mM Sodium fluoride, 10mM Sodium pyrophosphate decahydrate, 1mM Sodium orthovanadate, 1mM phenylmethylsulfonyl fluoride, 1% IGEPAL) containing protease inhibitor cocktails in ice and centrifuged at 10,000 x g for 5 min at 4°C. Details are further described in detail in the Supplementary Materials.

### Subjects for behavior

Animals were housed in a Techniplast cages (Techniplast, West Chester, PA, USA) with standard bedding, shredded brown paper, a cardboard tube, and a Nestlet square (Jonesville Corporation, Jonesville, MI). Cages were placed in ventilated racks at a temperature (20 C-22.2 C) and humidity (∼25%) controlled vivarium maintained on a 12:12 light-dark cycle. Animals were provided with standard food and water *ad libitum*. The mouse colony was maintained by breeding *Ube3a* deletion males (B6.129S7-*Ube3a*^tm1Alb^/J; Jackson Laboratory, Bar Harbor, ME; Stock No. 016590) with female C57BL/6J (B6J) mice to maintain a paternal transmission of the mutant allele. The subject animals were generated by breeding *Ube3a* deletion females with C57BL/6J (B6J) males, producing maternally inherited *Ube3a* deletion animals (*Ube3a*^mat−/pat+^; *Ube3a*^mat-/pat+;^ Angelman Syndrome model) and wildtype littermate controls (*Ube3a*^mat+/pat+^; mat+/pat+). Identification of mice was conducted labeling neonate paw at PND2-4 with non-toxic animal tattoo ink (Ketchum Manufacturing, Brockville, ON, Canada). Tissue for genotyping was collected using tails clipped at age PND5-7 (0.5 cm) following the UC Davis Institutional Animal Care and Use Committee (IACUC) policy regarding tissue collection. Genotyping was conducted with RED Extract-N-Amp (Sigma-Aldrich, St. Louis, MO) using primers R1965 5′ GCT CAA GGT TGT ATG CCT TGG TGC T 3′, WTF1966 5′ AGT TCT CAA GGT AAG CTG AGC TTG C 3′, and ASF1967 5′ TGC ATC GCA TTG TCT GAG TAG GTG TC 3′ for Ube3a. Mice were socially housed in groups of 2-4 by sex after being weaned at PND21. All procedures were approved by the Institutional Animal Care and Use Committee of UC Davis and Brown University and conducted in compliance with the National Institutes of Health Guide for the Care and Use of Laboratory Animals.

### Drug Administration of Syn3 and D-Syn3

Syn3 and D-Syn3 cyclic peptidomimetic compounds (WuXi AppTec) were dissolved in 1X phosphate buffered saline (PBS). Before testing, subjects were randomly assigned using a random number generator to receive Syn3, D-Syn3, or Vehicle (PBS 1X). Syn3 and D-Syn3 solutions were made fresh prior to every task. Syn3 and D-Syn3 were administered 2 hours prior to every behavior task, at a dose of 1 mg/kg via intraperitoneal (*i.p*.) injection, similar to other pharmacology studies in our laboratory [10,35–38].

### Behavioral Testing

To reduce carry-over effects from repeated behavioral testing, at least 24 hours were allowed to pass between the completion of one task, and the start of the next task in the order of the behavioral battery [5,7,10,39]. On days between behavioral tasks, animals were not injected. Assays were performed in the order of least to most stressful between the hours of 8:00AM PST and 7:00PM PST during the light phase. Group sizes were chosen based on experience and power analyses [40–48]. The experimenter was blind to genotype and treatment group. All surfaces of the testing apparatuses were cleaned using 70% ethanol. Mice were tested between 8-12 weeks of age and were assayed in the testing order: 1) open field locomotion for gross exploratory behavior; 2) DigiGait ™ for spatial and temporal indices of gait; 3) accelerating rotarod for motor coordination and motor learning; 4) short term novel object recognition; and 5) lethal dose (80 mg/kg, i.p.) pentylenetetrazol-induced seizures.

All behavior testing was conducted by an experimenter blind to genotype and treatment and both sexes were included. There were four treatment groups assessed 1) **WT** = control wildtype (*Ube3a*^mat+/pat+^) treated with vehicle (PBS 1X, i.p.) 2) **AS** = AS mouse model (*Ube3a*^mat-/pat+^) treated with vehicle (PBS 1X, i.p.) 3) **Syn3** = AS mouse model *Ube3a*^mat-/pat+^ treated with Syn3 (1 mg/kg, i.p.) and 4) **D-Syn3** = *Ube3a*^mat-/pat+^ mouse treated with D-Syn3 (1 mg/kg, i.p.).

### Open field

A novel open field arena was used to assess gross motor exploratory behavior, as previously described [5,10,49,50], for thirty minutes at ∼ 30 lux, described in the Supplementary Materials and shown in **Fig. S3**. Open field also required a two-way repeated measures ANOVA, comparing genotype and treatment over time.

### DigiGait ™

Metrics of gait were assessed using the DigiGait automated treadmill system and analysis software (Mouse Specifics Inc., Framingham, MA), as previously described [5,10], and described in detail in the Supplementary Materials, and data shown in **Fig. S4**.

### Accelerating rotarod

Motor coordination, balance, and motor learning was assessed using an Ugo-Basile accelerating rotarod (Stoelting Co, Wood Dale, IL), as described [7,10,35,51–53]. Subjects were placed on a rotating cylinder that starts at five revolutions per minute that accelerated from five to forty revolutions per minute over five minutes (300 seconds). For rotarod, repeated measures ANOVA for training day, was used to assess rotarod performance, analyzing motor coordination on Day 1 and motor learning by comparing Day 1 to Day 3, requiring a two-way ANOVA for training by genotype and treatment, followed by Holm-Sidak post hoc analysis.

### Novel object recognition (NOR)

The NOR assay was conducted in a 30-lux room in an arena (41 cm, 41 cm w x 30 cm h), described earlier, [7,10,37,51,52,54,55] and is described in the Supplementary Materials.

### Pentylenetetrazol-induced seizures

Pentylenetetrazol (PTZ) is a GABA_A_ receptor antagonist used to assess seizure susceptibility. Subjects were administered a lethal PTZ dose of 80 mg/kg i.p. and were immediately placed in a clean empty standard mouse cage. For a thirty-minute observation period, subjects were scored using indices of a modified Racine scale, as described [9,39,50]. Seizure parameters, following the induction of a behavioral seizure, were compared using a one-way ANOVA, followed by a Dunnett’s post hoc analysis, compared to vehicle treated WT, when p < 0.05. Survival curves were monitored and analyzed using a log-rank Mantel Cox test.

### Statistical Analysis

GraphPad Prism 9 (GraphPad Software, San Diego, CA) was used to perform statistical analysis and create graphs. Sex differences have not been observed, in this AS mouse model, using this tailored battery for AS rodent models, to date, and thus combined sexes were utilized to achieve ample power [5,7,10], and sex differences were not evaluated. Behavioral analysis passed normality distribution, using D’Agostino and Pearson tests. Data was collected using continuous variables, and thus, was analyzed via parametric analysis. Significance was based on an alpha of 0.05. Comparisons were between the WT and AStreated vehicle as well as AS treated with Syn3 or D-Syn3 to identify previously reported AS deficits and the effects of Syn3 and/or D-Syn3. Post hoc testing for multiple comparisons was carried out using Dunnet’s, Tukey’s, or Holm-Sidak’s multiple comparisons test.

## Results

### Discovery of novel cyclic TrkB-PSD-95 complex enhancing compounds of the peptidomimetic class

Our prior studies found that aberrant BDNF-TrkB-PSD-95 signaling contributes to the pathophysiology of AS [30,31]. As shown in the schematic in **Fig. 1A**, at excitatory synapses, the association of the postsynaptic scaffolding protein PSD-95 with TrkB is pivotal for BDNF-induced PLC-CaMKII and PI3K-Akt signaling, and in AS mice the BDNF-induced PSD-95-TrkB complex formation is impaired[31]. CN2097, a bridged cyclic peptide designed to bind the PDZ3 domain of PSD-95, facilitated PSD-95-TrkB complex formation to mitigate signaling deficits in an AS mouse model [31]. Although CN2097 shows a higher affinity for the PDZ3 domain (K_D_∼ 190nM) [32], further studies found that it also binds the PDZ1 domain [32]. Nevertheless, from this data, we inferred that selective binding to the PSD-95 PDZ3 domain plays a key role in promoting TrkB-PSD-95 complex formation and hypothesized that increasing affinity for the PSD-95 PDZ3 domain would increase efficacy. **Fig. 1B** shows the amino acid sequence of the novel cyclic-peptide Syn3, whose efficacy was evaluated in this study. As illustrated (**Fig. 1B**), in contrast to CN2097 that binds the PDZ3 domain, Syn3 binds the PDZ3 domain and an adjoining αC helix of PSD-95 to achieve bivalent binding that results in up to 5-fold stronger affinity (K_D_∼ 41nM), as reported previously by Naik et al. [32]. To increase peptide stability, we engineered analogs, D-Syn3, using dextrorotary (D)-amino acids that are less susceptible to proteolysis. D-Syn3 is identical in sequence, but Pro5 was substituted with the D-enantiomer (d-Proline) residue which was found to improve binding affinity by 2-fold, as well as substituting the entire arginine tag and cysteines with d-amino acids [32]. Following IP injection (1 mg/kg dose), microdialysis confirmed that Syn3 and D-Syn3 cross the blood-brain barrier (BBB) to reach a peak concentration in ∼90 mins and remain at detectable (≥ 0.1uM) concentrations for at least 7 h, with the D-Syn3 reaching a greater concentration (Cmax) (**Fig. 1C**). To test whether the Syn3 compounds can mitigate deficits in BDNF signaling resulting from decreased *Ube3A* expression, we tested the ability of D-Syn3 to reduce signaling deficits in human induced pluripotent stem cells induced neurons (iNs) by forced expression of Neurogenin-2 (NGN2) [33], in which Ube3A was knocked down as described previously [31]. As shown in Fig.1D, western blot analysis probed with phospho-specific antibodies demonstrates that Ube3A knockdown elicits deficits in the TrkB/PI3K/AKT/mTOR pathway (see also supplemental Fig.1 for the full set of data and quantification). The phosphorylation of Akt at regulatory residues Thr-308 and Ser-473 (S473, T308), an indicator of Akt kinase activation, was significantly decreased (siNC vs siUBE3A: 192.9%±9.5% vs 114.1%±2.8% for S473 and 141.3%±3.8% vs 100.3%±11.9% for T308; p<0.05; Fig.1D). Similarly, phosphorylation of the S6 ribosomal protein (Ser235/236), a downstream target of mTOR, was significantly decreased following Ube3A knockdown (siNC vs siUBE3A:167.4%±20.6% vs 92.5%±11%; p<0.05). Treatment with D-Syn3 at 0.1 uM, a ten-fold lower concentration compared to CN2097 reported to promote signaling [30,31], restored BDNF-induced Akt (S473, T308), and S6 ribosomal protein (Ser235/236) phosphorylation to levels indistinguishable from control iNs (**Fig.1D** and **Supplementary Fig. S1D** for quantification). As shown in **Supplementary Fig. S1**, further studies confirmed that treatment with D-Syn3 restored mTOR and CaMKII activity in siUbe3A iNs to levels indistinguishable from the level of control iNs (Fig. S1C for immunoblots and Fig. S1D for quantification). In iNs we sometimes observed that D-Syn3 alone, which normally acts only in the presence of BDNF could enhance TrkB signaling, reflected by increased Akt (S473, T308) and S6 ribosomal protein (Ser235/236) phosphorylation (**Fig.1D** and **Supplementary Fig. S1C, D**). This appears to be the result of basal levels of BDNF released by iNs, as this effect was abrogated by the TrkB inhibitor K252a, confirming that D-Syn3 is acting via TrkB in conjunction with endogenously released BDNF **(Supplementary Fig. S2).**

**Figure 1.**
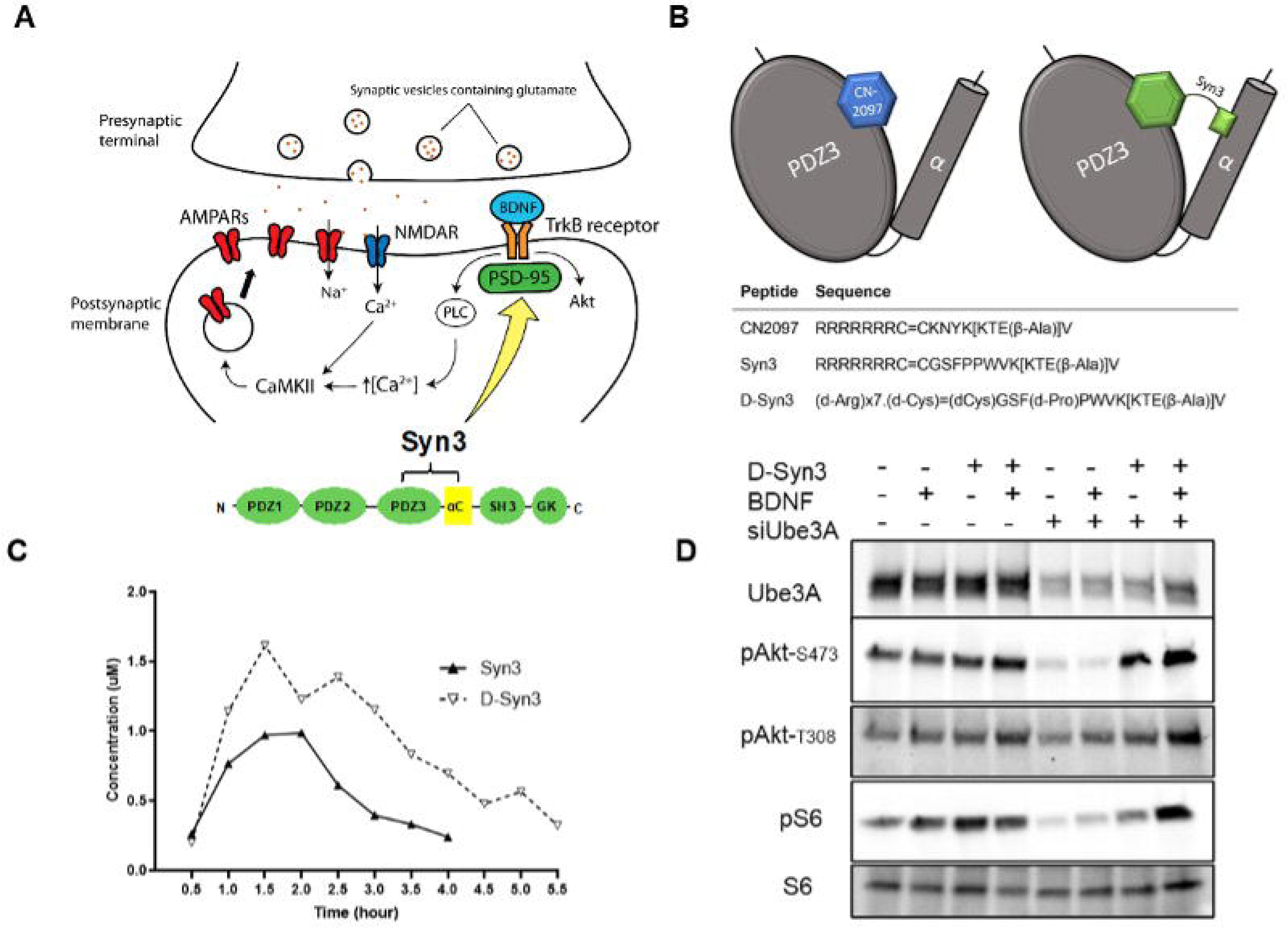
Targeting BDNF signaling. **(A)** A schematic showing the mechanism of action of Syn3 at the postsynaptic density. Syn3 binds PSD-95 to promote TrkB-PSD-95 association that facilitates PLC-CaMKII and PI3K-Akt signaling. **(B)** Schematic showing CN2097 binding to the PDZ3 domain whereas Syn3 bivalently binds the PDZ3 domain and αC helix of PSD-95. Shown are the amino acid sequences of CN2097, Syn3 and D-Syn3. All three peptidomimetics contain a cyclic moiety KTE(β-Ala)] V that is selective for the PDZ3 domain of PSD-95 and a polyarginine (R_7_) active-transport moiety to allow blood-brain barrier permeability. Syn3 and D-Syn3 contain additional PSD-95 αC helix-interacting residues derived from SynGAP. D-Syn3 was synthesized with D-amino acids: Pro5 substituted by d-Proline residue, and the entire arginine tag and cysteines. **(C)** Average microdialysis traces (n=3) of the mouse hippocampus after I.P. delivery of 1mg/kg Syn3 or D-Syn3 demonstrate that Syn3 compounds cross the BBB. **(D)** D-Syn3 rescues impaired TrkB signaling in siUBE3A iNs. Representative immunoblots of Akt-pThr308, pAkt-pSer473, S6-pSer235/236 in D-Syn3 treated siNC (control) and siUbe3A knockdown iNs (+) with or without (+/-) BDNF stimulation (see Fig. S1 for quantification of fold increase by BDNF).

### Syn3 and D-Syn3 rescue deficits in long term potentiation (LTP) in the Angelman Syndrome mouse model

Hippocampal long-term potentiation (LTP) is impaired in AS mice [8,56]. As reported in our previous study, the cyclic peptidomimetic CN2097 can restore LTP in AS mice at a dose of 10 mg/kg [31]. To examine the effects of the novel high affinity Syn3 compounds on synaptic plasticity, excitatory postsynaptic field potentials (fEPSPs) were recorded from the CA1 region, and LTP was saturated by repeated induction at Schaffer collateral afferents, as previously described [30,31]. Based on the in vitro signaling studies showing that D-Syn3 is effective at a ten-fold lower concentration compared to CN2097 (0.1uM D-Syn3 v 1uM CN2097; **Fig. 1** and **Supplemental Fig. S1**), and microdialysis showing that i.p injection of 1mg/kg Syn3 compounds remain detectable (≥ 0.1uM) in the brain for at least 7 h, a dose of 1mg/kg of Syn3 and D-Syn3 was chosen for these studies. To examine the effects of Syn3 compounds on synaptic plasticity, mice were injected with Syn3 or D-Syn3 (1mg/kg, i.p.) or vehicle (PBS 1X). Brain slices were prepared 2-3 hours after the injection and fEPSPs recorded from the CA1 region. HFS at Schaffer collateral afferents (2x 100 Hz for 1s separated by 20 s) elicited LTP in slices prepared from vehicle injected WT mice (183.9 ± 10.3%; **Fig. 2A**) but did not elicit significant LTP in slices prepared from vehicle injected AS mice, with a mean fEPSP slope change of 114.3 ± 6.3% (n=15 slices from 7 mice; **Fig. 2B**), which is compatible with previously reported results from AS mice [30,31]. Syn3 successfully restored LTP in AS brain slices to 205.4 ± 29.5% (n=11 slices from 5 mice, *p*<0.05; **Fig. 2C**) and D-Syn3 restored LTP to 187.1 ± 14.1% (n=20 slices from 11 mice, *p*<0.05; **Fig. 2D**). There was no significant difference in saturated LTP between Syn3 and D-Syn3 treatment (p=0.52). In addition, the amount of saturated LTP in Syn3 and D-Syn3 treated mice was comparable to WT slices [30].

**Fig. 2.**
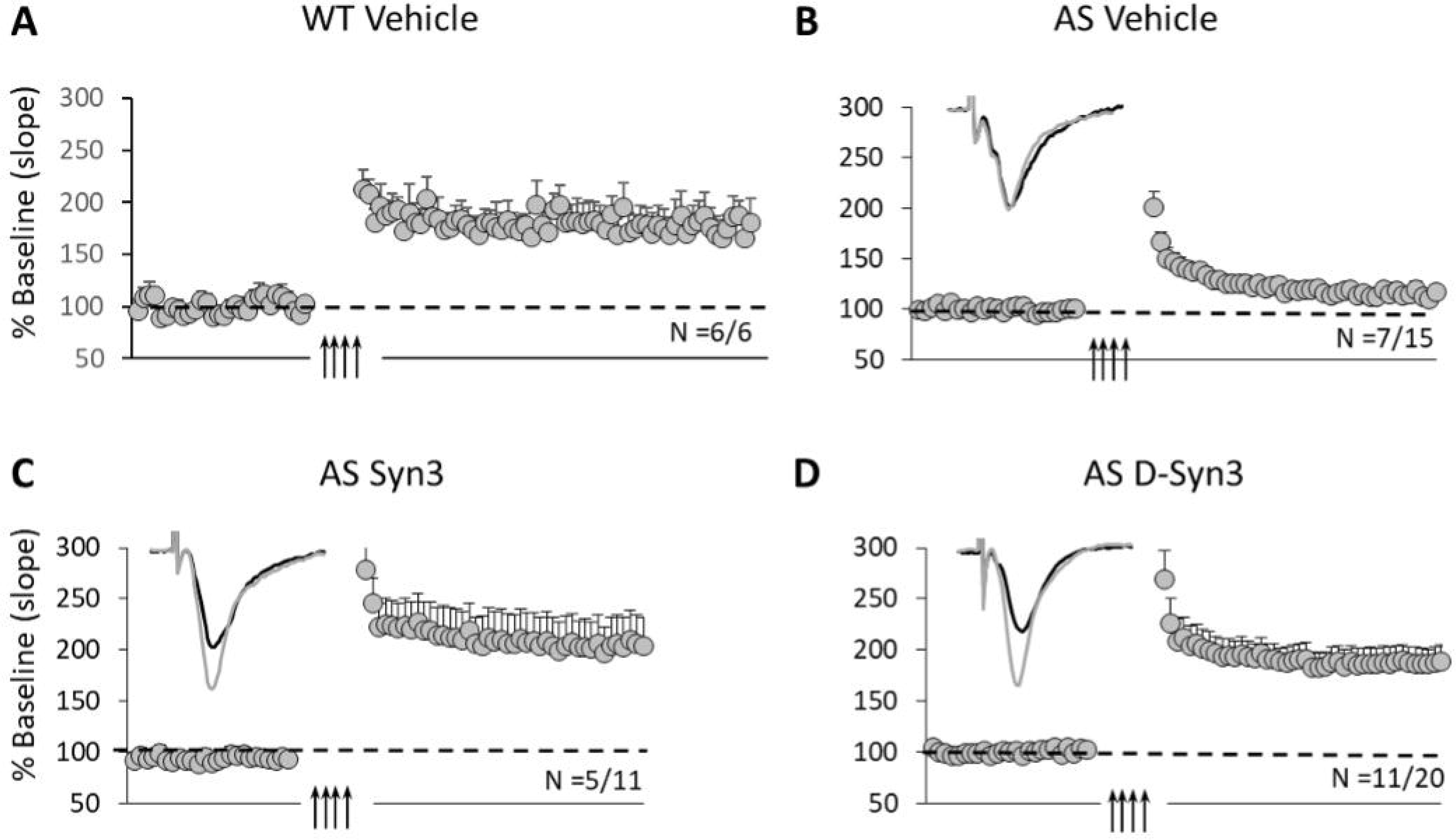
Syn3 and D-Syn3 restore synaptic plasticity in hippocampal slices of compound-injected AS mice. **(A-D)** Illustration of saturated LTP and representative traces (average of 10) before and after LTP saturation for all treatment conditions. fEPSP slopes were represented as percentage of baseline (Mean ± SEM). **(A)** LTP was observed in vehicle-injected WT mice (183.9±10.3). **(B)** AS mice treated with vehicle expressed no significant LTP (114.3±6.3%). **(C-D)** Syn3 treatment **(C)** and D-Syn3 treatment **(D)** of AS mice resulted in normal amounts of LTP (205.4±29.5% and 187.1±14.1% respectively). There was no significant difference in saturated LTP between the compounds (p=0.52).

### Syn3 and D-Syn3 improved seizure threshold and hyperexcitability in AS mice

To assess if Syn3 or D-Syn3 have antiseizure properties, we used a classic pentylenetetrazol (PTZ) seizure assay, administering a single i.p. injection of PTZ (80 mg/kg) and a modified Racine scale [57–60] to collect indices of seizure susceptibility [6,9,39,50,61–67]. Latency to myoclonic jerk trended toward exhibiting a faster onset but did not reach significance (**Fig. 3A**; F _(3, 56)_ = 1.447, NS). Therefore, we did not assess multiple comparisons. Compared to WT mice, AS mice exhibited reduced latency to generalized clonus seizure, tonic extension, and death (**Fig. 3B**; F _(3, 55)_ = 7.999, p < 0.0002; **Fig. 3C**; F _(3, 51)_ = 6.507, p < 0.0008; **Fig. 3D**; F _(3, 56)_ = 7.357, p < 0.0003).

**Figure 3.**
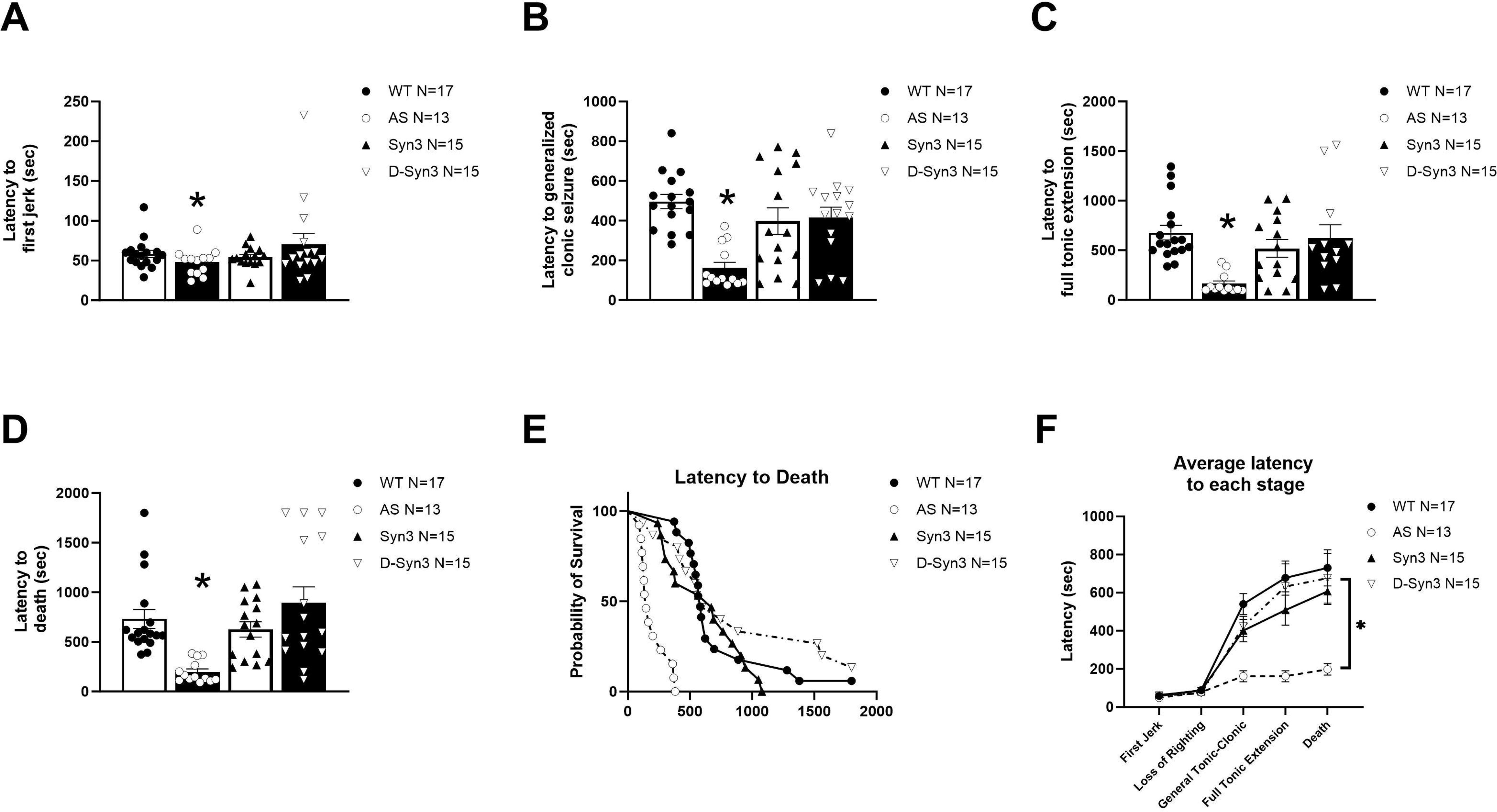
AS mice demonstrate increased seizure susceptibility across standard Racine scoring stages. Indices of seizure susceptibility were observed for 30 minutes following an i.p. injection of 80 mg/kg of pentylenetetrazole (PTZ). Syn3 and D-Syn3 lowered the heightened seizure susceptibility in *Ube3a*^mat-/pat+^ mice by multiple metrics. AS mice demonstrate reduced: **(A)** latency to first jerk, compared to WT, which is not observed following treatment with Syn3 and/or D-Syn3; **(B)** onset to generalized clonus, **(C)** latency to full tonic extension, **(D)** and latency to death. **(E)** The survival curve illustrates the difference in efficacy of Syn3 and D-Syn3 for each seizure stage, illustrating marginally improved effects from DSyn3K. **(F)** The illustration in Panel F highlights the entirety of the behavioral seizure in AS, WT, Syn3 and DSyn3K groups across the 30-minute period. One-way ANOVA and a Sidak’s multiple comparisons test. \**p<0.05* indicates when AS, Syn3 and D-Syn3 groups differ from the control WT group.

Notably, pre-treatment with Syn3 nor D-Syn3 in AS mice differed from WT (**Fig. 3B _Syn3_,** p = 0.3435; **Fig. 3B _D-Syn3_,** p = 0.5010) when assessing latencies to generalized clonus seizure. However, pre-treatment of Syn3 and D-Syn3 differed from vehicle treated AS (**Fig. 3B_Syn3_,** p < 0.0049; **Fig. 3B _D-Syn3_,** p < 0.0023), illustrating restoration and increased latency to generalized clonus seizure. Similarly, when assessing latencies to full tonic extension and death, pre-treatment of Syn3 nor D-Syn3 in AS mice differed from WT (**Fig. 3C _Syn3_,** p = 0.4136; **Fig. 3C _D-Syn3_,** p = 0.5010; **Fig. 3D _Syn3_,** p = 0.8117; **Fig. 3D _D-Syn3_,** p = 0.5443). However, pre-treatment of Syn3 and D-Syn3 in AS mice differed from vehicle treated AS (**Fig. 3C_Syn3_,** p < 0.0204; **Fig. 3C _D-Syn3_,** p < 0.0032; **Fig. 3D _Syn3_,** p < 0.0201; **Fig. 3D _D-Syn3_,** p < 0.0001) illustrating restoration and increased latency to full tonic extension and longer latencies to death. Survival curve analysis, shown in **Fig. 3E**, assayed with the log rank Mantel-Cox test revealed a Chi^2^ value of 60.45_df=3,_ p < 0.0001, illustrating that Syn3 and D-Syn3 lengthened the time to reach each of the Racine Score stages. AS mice treated with D-Syn3 had a 60% greater latency to reach full tonic extension compared to mice treated with Syn3. Moreover, both Syn3 and D-Syn3 had a significant influence on reducing seizure susceptibility across all seizure metrics observed in the 30-minute period (**Fig. 2F**). It appears as if D-Syn3 had a marginally greater effect on increasing latency to death and the last three of the five stages quantified in **Fig. 3F**.

### Syn3 and D-Syn3 rescued deficits in novel object recognition in AS mice, improving cognition

Novel object recognition (NOR) was performed to evaluate improvement in cognition, resulting from Syn3 or D-Syn3 pre-treatment. Vehicle treated WT mice spent more time investigating the novel object versus the familiar object, demonstrating intact recognition memory (**Fig. 4A**; t_(13)_ = 4.958, p < 0001), whereas vehicle treated AS mice did not (**Fig. 4A**; t_(12)_ = 1.205, p = 0.2515), as previously published [7,10]. Interestingly, like the WT, Syn3 and D-Syn3 treated AS mice also spent greater time investigating the novel object versus the familiar object, analyzed using within genotype paired t-tests (**Fig. 4A**; Syn3 t_(14)_ = 4.535, p < 0.0001; D-Syn3, t_(13)_ = 3.446, p < 0.0002). During the familiar phase, no group exhibited bias for the left or right side nor for either object (**Fig. 4B**).

**Figure 4.**
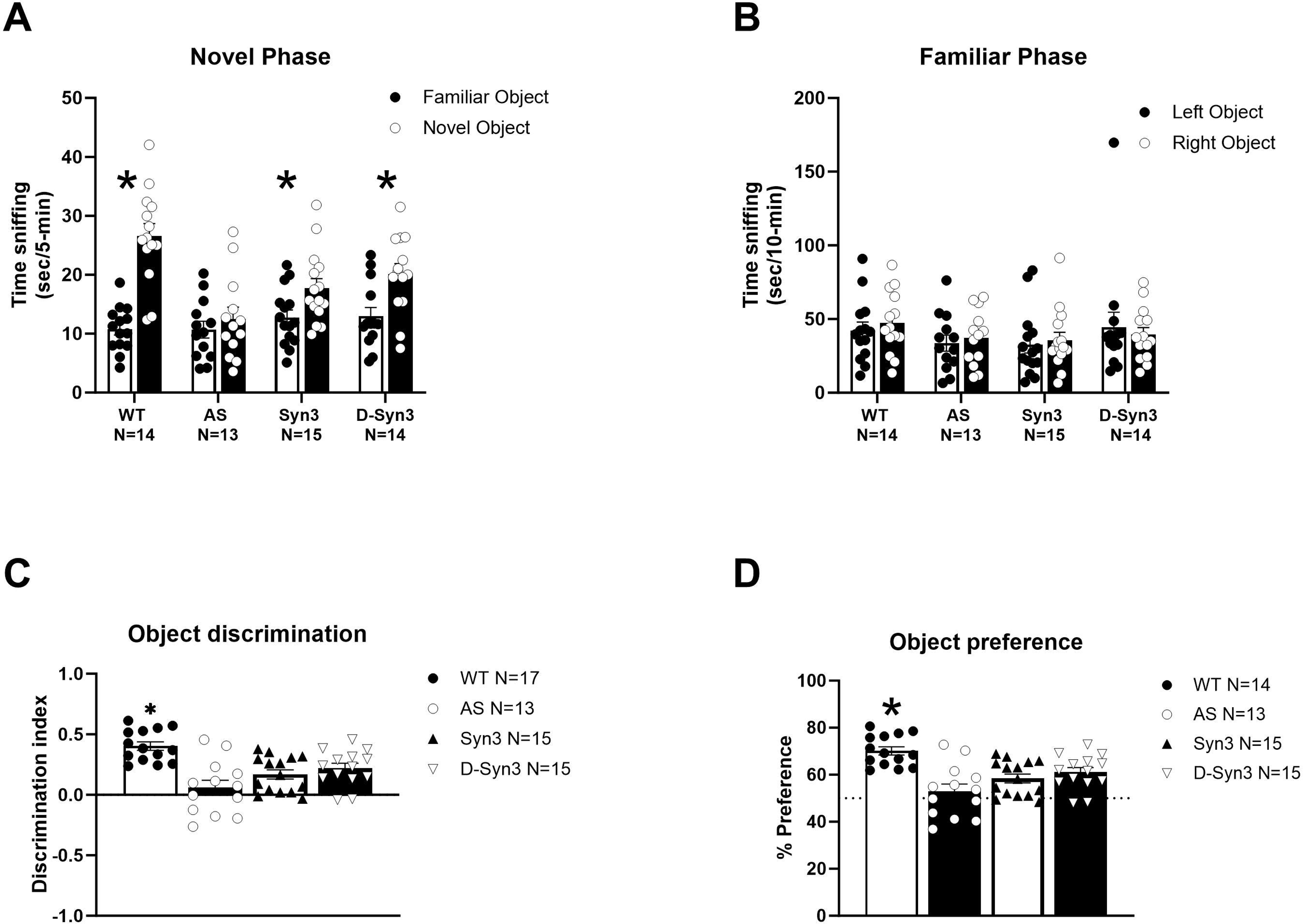
Subjects were evaluated for learning and memory abilities using the novel object recognition test. **(A)** AS mice did not spend more time with the novel object which illustrates a deficiency in recognition memory. Syn3 and D-Syn3 exhibited object recognition following a one-hour wait period, similar to WT. **(B)** During the familiar phase, none of the groups had a difference between the left and right object validating that neither object was preferred more the other. **(C)** Discrimination index reveals WT mice spend more time investigating the novel object compared to AS mice which is similar to findings in (**D)** novel object percent preference. \**p<0.05*, novel versus familiar, \**p<0.05* indicates when AS, Syn3 and D-Syn3 groups differ from the control WT group. Ordinary one-way ANOVA with Holm-Sidak multiple comparison test for *post-hoc* analysis.

Discrimination index analyzed treatment with Syn3 and D-Syn3 in AS mice, using one-way ANOVA (**Fig. 4C**; F _(3, 52)_ = 10.50, p < 0.0001), and discovered AS differed from WT (p < 0.0001). Syn3 and D-Syn3 also differed from WT, using Tukey’s post hoc analysis (**Fig. 4C**; Syn3, p = 0.0018; D-Syn3, p = 0.0253), but not vehicle treated AS (**Fig. 4C**; Syn3, p = 0.2994; D-Syn, p = 0.0590).

Preference ratio analyzed treatment with Syn3 and D-Syn3 in AS mice, using one-way ANOVA (**Fig. 4D**; F _(3, 52)_ = 10.50, p < 0.0001), and discovered AS differed from WT (p < 0.0001). Syn3 and D-Syn3 also differed from WT, using Tukey’s post hoc analysis (**Fig. 4D**; Syn3, p = 0.0018; D-Syn3, p = 0.0253), but not vehicle treated AS (**Fig. 4D**; Syn3, p = 0.2994; D-Syn3, p = 0.0590).

### D-Syn3, but not Syn3, rescued motor learning on the accelerating rotarod

Motor coordination and motor learning were measured using the accelerating rotarod. We observed improving performance over time, over the three training days (**Fig. 5A**; F _time (2, 108)_ = 77.30, p < 0.0001), genotype and treatment effects were also present, with significantly reduced latencies to fall compared to vehicle treated mice of the WT group (**Fig. 5A**; F_genotype/treatment (3, 56)_ = 24.05, p < 0.0001). We analyzed Days 1 and 3 to delineate Syn3 and D-Syn3’s effects on motor coordination versus motor learning in AS mice. When observing motor coordination of the groups on Day 1, a significant main effect between the groups was identified using two-way repeated measures ANOVA (**Fig. 5B**; F _(3, 56)_ = 24.17, p < 0.0001). On Day 1, following the Holm-Sidak post hoc analysis, Syn3 and D-Syn3 treated AS mice differed from WT (Syn3, p < 0.0001; D-Syn3, p = 0.0003). In contrast, on Day 1, Syn3 and D-Syn3 treated AS mice did not differ from vehicle treated AS (Syn3, p=0.3906, D-Syn3, p= 0.0790). On Day 3, our focused analysis on motor learning revealed a significant main effect between the groups was observed using two-way repeated measures ANOVA (**Fig. 5B**; F _(3, 56)_ = 19.06, p < 0.0001). Syn3 and D-Syn3 treated AS mice differed from WT (Syn3, p < 0.0001; D-Syn3, p < 0.0003) on Day 3. Additionally, on Day 3, Syn3 treated AS mice did not differ from vehicle treated AS, following Holm-Sidak post hoc analysis (Syn3, p = 0.0903). Notably, on Day 3, D-Syn3 treated AS mice significantly differed from vehicle treated AS mice (D-Syn3, p = 0.0124), suggesting an improvement in AS motor learning by D-Syn3.

**Figure 5.**
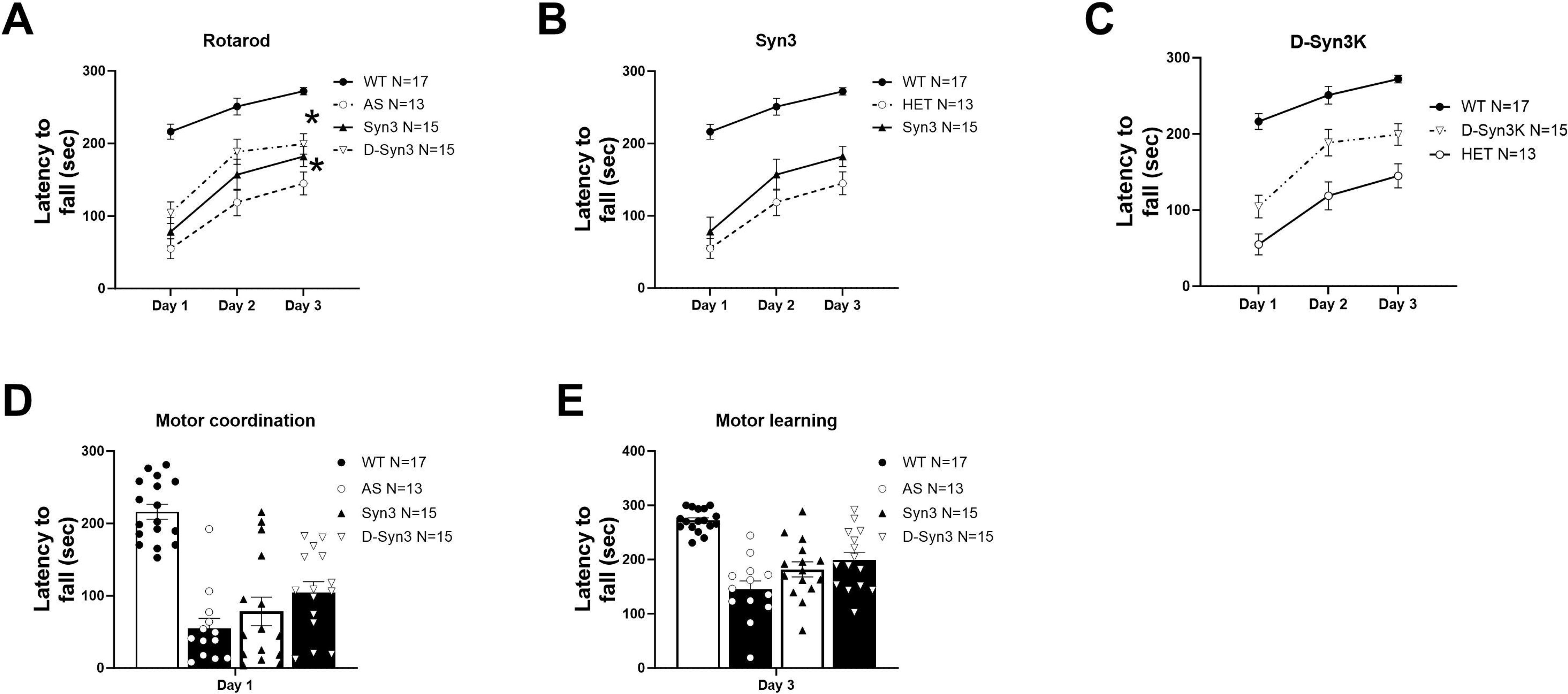
Accelerating rotarod was used to evaluate motor coordination and motor learning. **(A)** AS mice had a lower latency to fall compared to WT mice across all three days**. (B)** Syn3 did not rescue motor coordination nor motor learning in AS, yet trends for improvement were observable. **(C)** D-Syn3 exhibited improvements in AS mice, compared to WT, as the task paradigm was learned and repeated across days. To dissect the effect of time and test-retest of Syn3 and D-Syn3 in AS mice on the RR assay. We illustrate Day 1 **(D)** and Day 3 **(E)** performance, individually. Data are expressed as mean +/- SEM. \**p<0.05* indicates when AS, Syn3 and D-Syn3 groups differ from the control WT group. Performance on individual days was evaluated using a two-way RM ANOVA, followed by Holm-Sidak post hoc analysis.

## Discussion

BDNF has a well-established role as a regulator of synaptic plasticity, dendritic outgrowth, dendritic spine formation and maintenance. Reduced levels of BDNF signaling have been reliably observed in the AS mouse model [68,69]. Previously, our team reported BDNF-induced signaling requires TrkB association with PSD-95, and disruption of this association, in the AS mouse model, which likely underlies AS deficits in learning and memory [30,31] and elevated seizure and spike train phenotypes. In combination, these data suggest that drugs tailored to enhance the BDNF-TrkB-PSD-95 coupling would be an innovative approach for the treatment of AS and potentially a variety of NDDs.

Currently, PSD-95 targeted drugs have focused on the PDZ domains, yet developing a drug with high affinity and PDZ domain subunit specificity remains challenging, with three PDZ domains exhibiting high homology and a shallow binding pocket [32]. Here, we investigated a new class of cyclic-peptidomimetic ligands, Syn3 compounds, that bind the PDZ3 domain of PSD-95 with high affinity to enhance BDNF-TrkB signaling [32]. BDNF signaling studies in human IPSC iNs confirmed that the Syn3 compounds are bioactive at a 10-fold lower concentration, compared to our previously described CN2097 compound [31]. In iNs, we found that *Ube3A* knockdown resulted in decreased PI3K/Akt/mTORC1 signaling and impaired αCaMKII activity due to increased inhibitory phosphorylation of Thr305/306 [70,71](Fig.S1; *p*<0.05). Treatment with the Syn3 analog D-Syn3 reduced the excessive CaMKII autoinhibitory phosphorylation and restored BDNF-induced Akt, mTOR and S6 ribosomal protein phosphorylation to levels indistinguishable from control iNs (**Fig.1D and Fig. S1**). In alignment with the signaling studies, *i.p.* injection of AS mice with Syn3 compounds at a 10-fold lower dose compared to CN2097 (1mg/kg versus 10 mg/kg) restored LTP. Additionally, we observed the binding of the Syn3 drug to the PDZ3 domain of PSD-95 is inversely proportional to its potency, which illustrates Syn3 compounds target PSD-95 *in vivo*. Furthermore, when evaluating Syn3 and D-Syn3 in translationally relevant behavioral assays, we observed improvements in seizure threshold, cognitive deficits, and LTP dysfunction.

Seizures are predominant in AS (∼80%), yet classic antiepileptic drugs remain ineffective. Syn3 compounds exhibited remarkable *in vivo* seizure efficacy and improvement in the AS mouse model following seizure induction by the GABA_A_ receptor antagonist PTZ, a common chemically induced seizure model. PTZ-induced convulsions were used to gauge susceptibility to primary generalized seizures and as a gross approximation of excitation-inhibitory balance, which has been hypothesized to underlie AS and other genetic NDDs. Systemic administration of Syn3 and D-Syn3 reduced the seizure threshold across behavioral indices of the modified Racine scoring, including generalized clonic-tonic seizure, tonic extension, and death [59,60]. Current studies testing potential therapeutics in preclinical AS mouse models commonly use audiogenic paradigms and/or flurothyl kindling [62,63,66,72,73] or AS mice on a 129SvEv background, which are basally seizure susceptible because of the greater frequency and increased severity of callosal agenesis and hippocampal commissure defects [6,11,60]. Moreover, this report is the first of its kind as our findings were generated using the maternal *Ube3a* mutation mice on a pure C57BL/6J background strain, [60,74–76], over the seizure susceptible, 129SvEv. Notably, the restored seizure threshold to WT levels following treatment of Syn3 and D-Syn3 illustrates its promise as a potential antiepileptic drug.

Syn3 and D-Syn3 improved cognition suggesting that improvements in behavioral phenotypes are possible beyond a critical period. Using the novel object recognition assay as a measure of learning and memory, both Syn3 and D-Syn3 treated AS mice exhibited improvements comparable to WT. In addition, rotarod performance across three days demonstrated a rescue in motor learning by D-Syn3 that was more effective than Syn3. The improved efficacy of D-Syn3 in motor learning could be the result of D-Syn3 reaching a higher Cmax or remaining at therapeutic doses in the brain for longer times, as shown by microdialysis (**Fig. 1C**). Neither Syn3 nor D-Syn3 improved gross exploratory locomotion deficits, motor coordination, gait impairments, behavioral impairments that have previously been well documented in the AS rodent models, which highlight the need for further pharmacokinetic investigation (**Fig. S3, Fig. S4**). The current work did not explore alternative dosing paradigms, routes of administration, or post-treatment intervals, which could increase efficacy in a broader number of domains. Although gross and fine motor deficits did not improve with acute systemic treatment of Syn3 and D-Syn3, chronic treatment may improve these outcomes. Additionally, this study was conducted in adult mice from 8-12 weeks of age. Given that the behavioral deficits of AS emerge early in life, future studies could include a range of ages assessed to identify critical windows of effective treatment.

Moreover, aberrant BDNF signaling has been proposed to contribute to the pathophysiology of a variety of neuropsychiatric disorders, especially those with a substantial cognitive component. In neuropsychiatry, the discovery of cognitive enhancers or drugs that would minimize impairments in learning and memory for ID disorders or the progressive decline of cognition in neurodegenerative disorders would be transformative in preclinical neuropharmacology. Furthermore, single gene disorders, involving synaptic proteins that interact with PSD-95 [77,78] are plentiful, such as Phelan-McDermid Syndrome (PMS) involving a loss of *SHANK3* and SynGAP1-R Intellectual Disabilities (SRID), resulting from a loss of a single copy of *SYNGAP1*. Future studies should assay Syn3 and D-Syn3 in PMS or SRID preclinical models given that our compounds are closely related to binding cascades involving SynGAP1 and Shank3 [79]. In summary, Syn3 and D-Syn3 effectively improve seizure threshold and cognitive deficits in a mouse model of AS. Our findings emphasize the promising utility of Syn3 and D-Syn3 as a potential therapeutic for these behavioral domains. These findings highlight the necessity for future preclinical studies in other animal models to assess Syn3 and D-Syn3 as a therapeutic for AS and additional NDDs.

## Acknowledgements

We thank the Angelman Syndrome community and the UC Davis MIND Institute for supporting this research. We also thank Dr. Nathaniel Hodgson (IDDRC Animal Behavior and Physiology Core) for performing microdialysis and Rory (Dallon) Martin for exceptional husbandry and attention to the mouse colonies of the Silverman Laboratory and the IDDRC mouse behavioral core at the MIND Institute at UC Davis School of Medicine.

## Author information

These authors jointly supervised this work: Jill L. Silverman, John Marshall.

### Contributions

JLS and JM designed and funded the study. EZH performed the behavioral experiments and subsequent analyses. XY and YAH conducted the biochemistry and signaling studies in the iN cells. MSP conducted the LTP analysis. MN completed the structural biology analysis. JM and JLS supervised the study and interpretations of data. EZH, JM, and JLS drafted the initial manuscript. All authors included valuable comments and edits to the manuscript. All authors read and approved the final manuscript.

## Ethics declarations

