## Supplementary Materials for "Peptidomimetic inhibitors targeting TrkB/PSD-95 signaling improves cognition and seizure outcomes in an Angelman Syndrome mouse model"

**Supplementary Methods and Materials**

***Pluripotent stem cell induced neurons (iNs)***

iNs were generated following previous protocol by forced expression of Neurogenin-2 (NGN2) with kolf2.J1 stem cell lines [54]. The iNs were cultured in neural basal medium (Thermo Fisher Scientific 10888022) supplemented by N2 (Thermo Fisher Scientific 17502001) and B27 (Thermo Fisher Scientific 17504044). SiRNA targeting UBE3A (SiUBE3A), and relative negative control (SiNC) were ordered from Thermo Fisher Scientific (Thermo Fisher Scientific 4390815 and AM4615) and used at 100nM final concentration at ~DIV8 and incubated for 2-3 days before drug treatments.

***Western blot***

All samples were harvested in cell lysis buffer (1% SDS, 0.5% Deoxcholate, 50 mM Sodium phosphate diabasic heptahydrate, 150 mM Sodium chloride, 2mM EDTA, 50 mM Sodium fluoride, 10mM Sodium pyrophosphate decahydrate, 1mM Sodium orthovanadate, 1mM phenylmethylsulfonyl fluoride, 1% IGEPAL) containing protease inhibitor cocktails in ice and centrifuged at 10,000 x g for 5 min at 4°C. The supernatant was collected, protein concentration was measured by BCA protein assay, and further mixed with 4x Lamella sample buffer. After 10minutes boiling, samples were subjected to SDS-PAGE gel after brief spin down and transferred with Bio-Rad trans-bot system. Primary Antibodies (Ms-E6AP and Ms-Tubulin: Santa Cruz technology sc-166689 and sc-166729; Ms-GAPDH Proteintech #60004-1-IG; Rb-βCatenin, Rb-GSK3β-pSer9, and Ms-GSK3β Cell Signaling Technology #8480s, #9336s, and #9832S) were incubated overnight at 4 degrees Celsius after blocking in TBST buffers. Membranes were developed with HRP-conjugated secondary Antibodies (Cell Signaling Technology 7074S and 7076S) and filmed under iBright Imaging Systems. Band intensities for target proteins were measured by the ImageJ system (https://imagej.net/ij/ ).

***Electrophysiology in hippocampal slices to assess deficits in long term potentiation in AS mice and the effects of Syn3 and D-Syn3.***

Mice were deeply anesthetized (ketamine), decapitated and their brains quickly removed and immersed in oxygenated (95% O2/5% CO2) artificial cerebrospinal fluid (ACSF) containing (in mM): 126 NaCl, 3 KCl, 1.25 NaH2PO4, 1 MgSO4, 2 CaCl2, 26 NaHCO3, 10 glucose. Coronal brain slices (400 µm) containing the dorsal hippocampus and adjoining cortex were sectioned using a Vibratome, transferred to a temperature controlled (34±0.5°C) interface chamber and perfused with oxygenated ACSF at 1-2 ml/min. Brain slices were allowed to recover for at least one hour before recordings were started.

Extracellular postsynaptic field potentials (fEPSPs) were recorded using Borosilicate glass microelectrodes (resistance <1 MΩ) placed in CA1 stratum radiatum. Synaptic responses were elicited by stimulation of the Schaffer Collaterals with pulses of 0.2 ms duration at 0.03 Hz using concentric bipolar electrode (FHC Inc). Responses were amplified (AxoClamp2B, MolDevice and EX1, Dagan) and digitized at 10 kHz. Igor Pro software (Wave Metrics) and Neuromatic was used for data acquisition and analysis. The stimulation intensity eliciting 50% of maximum fEPSP amplitude was used for baseline recordings. Following 20-25 min stable baseline recordings LTP was induced by high frequency stimulation (2xHFS for 1s at 100 Hz separated by 20 s) as many times as necessary to saturate responses. fEPSP slopes were measured before and after LTP saturation and values expressed as percentage of baseline ± SEM. Paired two-tailed t-tests were used for statistical analysis.

***Open Field***

A novel open field arena was used to assess gross motor activity and exploratory behavior. Individual subjects were placed within a novel open arena (40 cm length x 40 cm width x 30.5 cm height) for thirty minutes at ~ 30 lux. VersaMax Animal Activity Monitoring System (AccuScan Instruments, Columbus, OH) was used to automatically detect photocell beam breaks to measure horizontal activity, total activity, vertical activity, time spent in the center, and total distance traversed. Data were analyzed using two-way repeated-measures ANOVA with genotype and treatment as between-group factors. Analysis of summed data was conducted using an ordinary one-way ANOVA with a Holm-Sidak’s multiple comparison test.

***DigiGait***

Metrics of gait were assessed using the DigiGait automated treadmill system and analysis software (Mouse Specifics Inc., Framingham, MA). Non-toxic red food coloring was used to paint the paws to minimize conflict with dark-green paw tattoos in DigiGait analysis. Before running the mice, the subjects were placed into Perspex walking treadmill to habituate. Consecutive strides at a belt speed of twenty centimeters per second for three-six seconds were recorded for each subject. The treadmill was cleaned with 70% ethanol after each run. Further gait analysis was conducted by an experimenter blinded to genotype and treatment. Data were analyzed per limb using two-way ANOVA with genotype and treatment as between-group factors. A post-hoc Holm Sidak multiple comparison test was used to assess performance between genotypes and treatment for each limb was conducted.

***Novel Object Recognition***

EthoVision XT video tracking software (Noldus Information Technology) was used to measure time spent investigating each object. Manual scoring by a trained observer blinded to genotype and treatment group was used to validate the automated scoring. Time spent sniffing the object when the nose was within 2 cm of the object and the placed toward the object was qualified as object investigation. If mice did not spend at least 5 seconds sniffing objects during the familiarization phase, they were moved from analysis. Performance was further evaluated using the discrimination index and percent preference. The discrimination index was calculated by dividing the difference in investigation time between the novel and familiar object by the total amount of exploration of both objects. Preference for an object was calculated as time spent sniffing the novel object compared to the total time sniffing both objects. Greater than fifty percent represents intact recognition whereas fifty percent lack of preference spending equal time investigating the novel and familiar object. Recognition memory was quantified as spending statistically significant more time investigating the novel object compared to the familiar object. For novel object recognition, object investigation was defined as time spent sniffing the object when the nose was oriented toward the object and the nose–object distance was 2-cm or less. Recognition memory was defined as spending significantly more time sniffing the novel object compared to the familiar object determined within genotype and treatment, using a paired t-test. Total time spent sniffing both objects was used as a measure of general exploration. Time spent sniffing two identical objects during the familiarization phase confirmed the lack of an innate side bias. Objects used were plastic toys: a small soft plastic orange safety cone and a hard plastic magnetic cone with ribbed sides. Analysis using discrimination index and preference ratio used a one-way ANOVA followed by Tukey’s post hoc analysis, which compares all groups to each other group.

**Supplemental Figures**

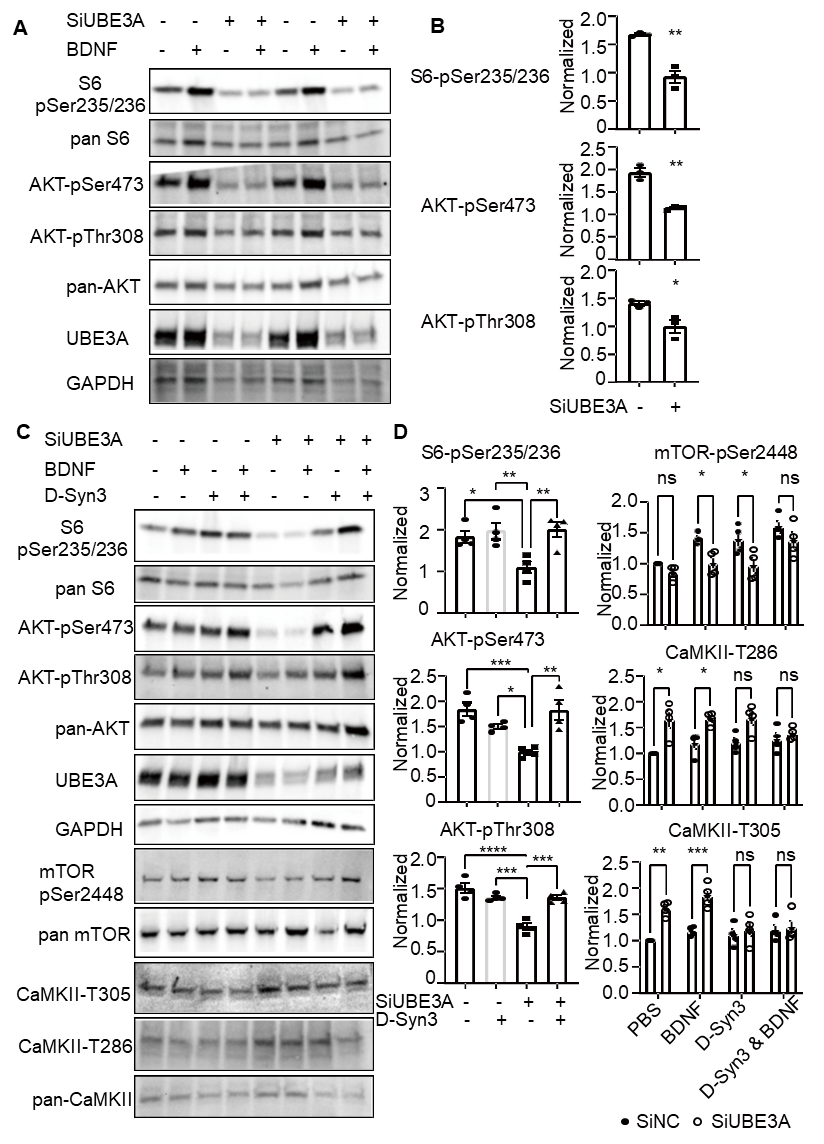

**Supplemental Figure S1.** **Impaired TrkB-BDNF signaling in siUBE3A iNs. Ube3A knockdown by siRNA reached over 90% at the protein level resulting in a significantly reduced BDNF-induced phosphorylation of the PI3K/Akt/mTORC1 signaling pathway. A**. Representative blots of S6-pSer235/236, AKT-pThr308, pAKT-pSer473 in siNC (control) and SiUBE3A iNs with or without (+/-) BDNF stimulation. **B**. Quantification of fold increase by BDNF; n=3 biological repeats for each condition. Unpaired T test, * p<0.05, ** p<0.01.  **C**. Representative blots of AKT-pThr308, pAKT-pSer473, mTOR-pSer2448, S6-pSer235/236, CaMKII-pT286 and CaMKII-p305 in D-Syn3 treated siNC and SiUBE3A iNs with or without (+/-) BDNF stimulation. We observed a decrease in mTOR phosphorylation at serine 2448 (Ser2448) (siNC vs SiUBE3A: 140.6%±4.4% vs 100.3%±9.9%). Ube3A knockdown iNs exhibited significantly increased (160.46%±6.99%, normalized SiUbe3a to siNC) αCaMKII at Thr305/306 phosphorylation and an increase (166.5%±16.7%, normalized siUbe3a to siNC) in Thr286 aCaMKII autophosphorylation (*p*<0.05). **D**. Quantification of fold increase by BDNF; n=4 biological repeats for each condition. One-way ANOVA with post-hoc analysis, * p<0.05, ** p<0.01, *** p<0.001, or ****P<0.0001.

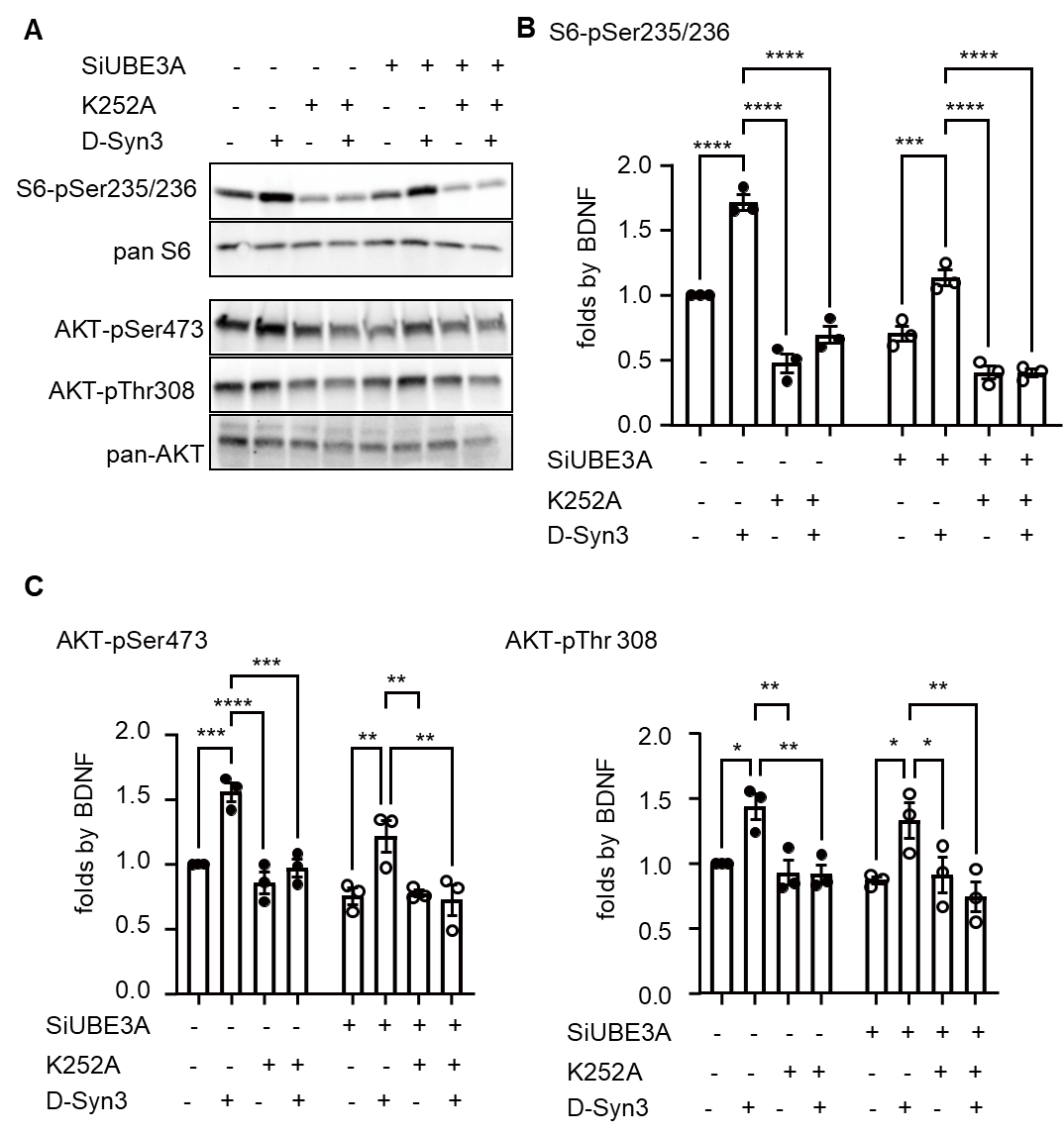

**Supplemental Figure S2.  D-Syn3 activates TrkB-BDNF signaling in SiUBE3A iNs. A**. Representative blots of S6-pSer235/236, AKT-pThr308, pAKT-pSer473 in D-Syn3 treated siNC and siUBE3A iNs, with or without K252A pre-incubation. **B-C**. Quantification of fold increase by D-Syn3. n=3 biological repeats for each condition. Two-way ANOVA with post-hoc analysis, * p<0.05, ** p<0.01, *** p<0.001, or ****P<0.0001.

  
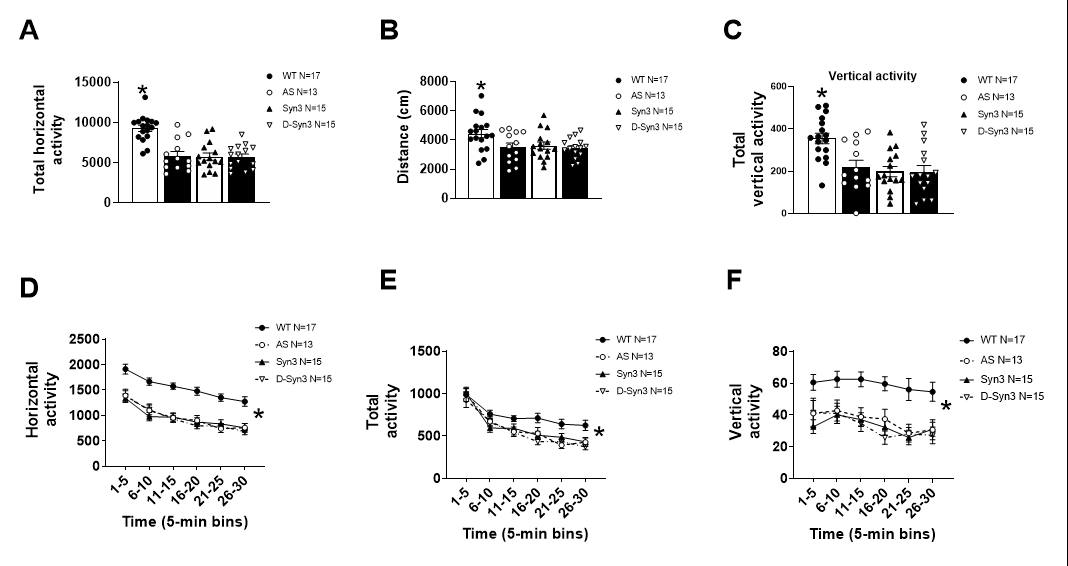

**Supplemental Figure S3**. **Syn3 and D-Syn3 did not improve novel exploratory locomotion in AS mice, which are substantially impaired, as previously published.** **(A)** Horizontal activity, **(B)** total activity, and **(C)** vertical activity in an open field assay were reduced in ***AS*** mice compared to **WT**. Syn3 and D-Syn3 had no effect on **(D)** horizontal activity, **(E)** total activity, or **(F)** vertical activity throughout the 30-minute exploration time. Ordinary one-way ANOVA with a Holm-Sidak’s multiple comparison test *p<0.05.

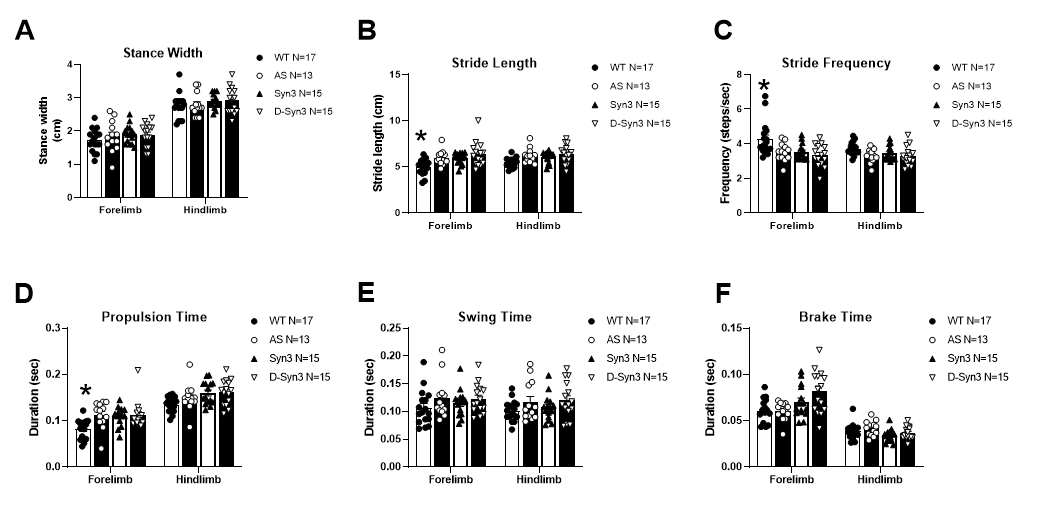

**Supplemental Figure S4. Syn3 and D-Syn3 did not affect spatial and temporal metrics of gait.** **(A)** There were no differences in stance width in AS mice compared to WT nor any effects of treatment. **(B)** AS mice had longer strides in the forelimb compared to WT, whereas **(C)** stride frequency was less compared to WT. **(D)** Propulsion duration as elevated in AS mice in the forelimb, however there was no difference in **(E)** swing duration or **(F)** brake duration between limb or genotype. Two-way ANOVA with a Holm-Sidak’s multiple comparison test *p<0.05.
